# Generation of a hybrid *App*^NL-G-F/NL-G-F^×*Thy1*-GCaMP6s^+/-^ Alzheimer disease mouse mitigates the behavioral and hippocampal encoding deficits of *APP* knock-in mutations of *App*^NL-G-F/NL-G-F^ mice

**DOI:** 10.1101/2022.11.18.517152

**Authors:** Samsoon Inayat, Brendan B. McAllister, HaoRan Chang, Sean G. Lacoursiere, Ian Q. Whishaw, Robert J. Sutherland, Majid H. Mohajerani

**Author notes:** Equal contribution (Samsoon Inayat, Brendan B. McAllister). Corresponding authors: Samsoon Inayat, Majid Mohajerani.

## Abstract

In contrast to most transgenic mouse models of Alzheimer disease (AD), knock-in mice expressing familial AD-linked mutations of the amyloid precursor protein (*App*) gene exhibit stereotypical age-dependent amyloid beta (Aβ) pathology and cognitive impairment without physiologically unrealistic *App* overexpression. This study investigated the effect of familial AD-linked *App* mutations on hippocampal CA1 neuronal activity and function. To enable calcium imaging of neuronal activity, *App^NL-G-F/NL-G-F^* knock-in (APPki) mice were crossed with *Thy1*-GCaMP6s^+/-^ (C-TG) mice to generate *App^NL-G-F/NL-G-F^*×*Thy1*-GCaMP6s^+/-^ (A-TG) mice, which were characterized at 12 months of age. A-TG mice exhibited Aβ pathology in the hippocampus. In several configurations of an air-induced running task, A-TG mice and C-TG mice were equally successful in learning to run or to stay immobile. In the Morris water place test, A-TG mice were impaired, but learned the task. Comparisons of hippocampal CA1 neuronal activity in the air-induced running task showed that A-TG mice displayed neuronal hypoactivity both during movement and immobility. A-TG mice and C-TG CA1 neuronal encoding of distance or time in the air induced running task were not different. These results suggest that knock-in of familial AD-linked mutations in A-TG mice results in Aβ pathology, neuronal hypoactivity, and cognitive impairment without severely affecting CA1 neuronal encoding. In comparison to APPki mice, A-TG mice had less severe AD-like memory impairments at 12 months of age (Saito et al., 2014; Mehla et al., 2019), suggesting that the disease onset was delayed in A-TG mice. The effect of *APP* mutations may have been mitigated through genetic mechanisms when APPKi mice were crossed with C-TG mice.

## INTRODUCTION

Alzheimer disease (AD) is a progressive neurodegenerative disorder causing life-altering behavioral and cognitive impairment, with no known cure. The symptoms of AD begin between 30 and 80 years of age, with a complex etiology governed by genetics, age, and environmental factors (McDonald et al., 2010; Carreiras et al., 2013, 2013). More than 200 mouse models have been generated to study the etiology and pathophysiology of AD, but no model accounts for the disease at all physiological levels, from genes to behavior (Alzforum, http://www.alzforum.org). At the cellular level, the study of neuronal activity and function is conducted to discover novel biomarkers (Busche and Konnerth, 2015; Stargardt et al., 2015) and to address how AD-associated gene mutations and pathological processes affect behavior (Cacucci et al., 2008a; Zhao et al., 2014a; Cayzac et al., 2015a; Busche and Konnerth, 2016; Mably et al., 2017a; Jun et al., 2020; Prince et al., 2021; Takamura et al., 2021), with the goal of elaborating therapeutic strategies (Busche et al., 2008, 2015; Iaccarino et al., 2016). The present study describes hippocampal neuronal function, focusing on distance and time encoding in a novel air-induced running task, in a second-generation AD mouse model, the *App^NL-G-F/NL-G-F^* knock-in (APPki) strain, which exhibits AD-like etiology and pathology (Saito et al., 2014).

Previous studies have described impaired neuronal function and impaired spatial behavior in transgenic AD mouse models that exhibit accumulation of amyloid beta (Aβ) in the brain, a hallmark pathological marker of AD (Busche et al., 2008; Cacucci et al., 2008b; Saito et al., 2014; Zhao et al., 2014b; Busche and Konnerth, 2015, 2016; Cayzac et al., 2015b; Mably et al., 2017b; Prince et al., 2021). A limitation of many of these transgenic models is that they overexpress amyloid precursor protein (*App*), from which Aβ is cleaved. *App* overexpression leads to the overproduction and accumulation of Aβ. This leads to increases in the production of neurotoxic *App* cleavage products that are not overproduced in human AD (Saito et al., 2014). To overcome the limitation of *App* overexpression, APPki mice have been generated, which express *App* with a humanized Aβ region and three mutations associated with familial AD (Saito et al., 2014). With age, these mice develop extracellular deposits of insoluble Aβ and display impairments in spatial navigation, object recognition, and cued and contextual fear memory and so more closely model human AD (Saito et al., 2014; Mehla et al., 2019).

Two studies have characterized spatial-behavior-related neuronal activity in APPki mice by examining the functional properties of place and time cells. In one study, hippocampal CA1 place cells of 7-13 months old APPki mice were found to be mildly altered, and their remapping properties were severely affected (Jun et al., 2020). No difference in the activity levels of neurons was observed. In the second study, hybrid *App^NL-G-F/NL-G-F^*/*Thy1*-G-CaMP7-T2A-DsRed2^+/-^ mice were generated from APPki mice to facilitate calcium imaging. For these mice, a reduction in active place cells and their specificity was found when comparing 7-month-old with 4-month-old mice, but no impairment was seen in behavior or in time cells (Takamura et al., 2021). Furthermore, cells near Aβ plaques were hyperactive. These conflicting results in relation to place cell function and neuronal activity levels could be attributed to the genetic strain (APPki vs. hybrid), the behavioral paradigm (freely moving on a linear track vs. head-fixed on a virtual reality based linear track), or to the method of recording neuronal activity (electrophysiology vs. calcium imaging). The present study was undertaken to clarify how *App* knock-in mutations and the formation of Aβ plaques affect spatial and temporal neuronal encoding in APPki mice and hybrid variants.

In the present study, *App^NL-G-F/NL-G-F^*×*Thy1*-GCaMP6s^+/-^ (A-TG) mice were generated by crossing the APPki strain with the *Thy1*-GCaMP6s^+/-^ strain, which expresses the genetically-encoded calcium indicator GCaMP6s under the control of the *Thy1* promotor (Dana et al., 2014). This allowed for calcium imaging of neuronal activity in a second-generation AD mouse model (Fig. 1A). *Thy1*-GCaMP6s^+/-^ (C-TG) mice were used as controls. The behavior of head-fixed mice was controlled by having them run a fixed distance and then stop on a conveyor belt task in response to an air stream. Neuronal encoding of event-related distance and time in hippocampal CA1 neurons was examined as a surrogate for place and time cell assessment. Spatial learning in the mice was assessed in the Morris place task. It was hypothesized that, at 12 months of age, A-TG mice would exhibit abundant Aβ plaques and disrupted event-related distance and time behavior and neuronal encoding, as well as impaired spatial navigation ability, in accordance with previous reports on APPki mice (Saito et al., 2014; Mehla et al., 2019; Jun et al., 2020) and a variant (Takamura et al., 2021). As will be described, the A-TG crossed strain exhibited no change in motor learning or CA1 neuronal encoding at 12 months of age as spatial behavioral impairments began to emerge, despite a lower average neuronal firing rate and numerous Aβ plaques in the hippocampus (Fig. 1B-E).

**Figure 1.**
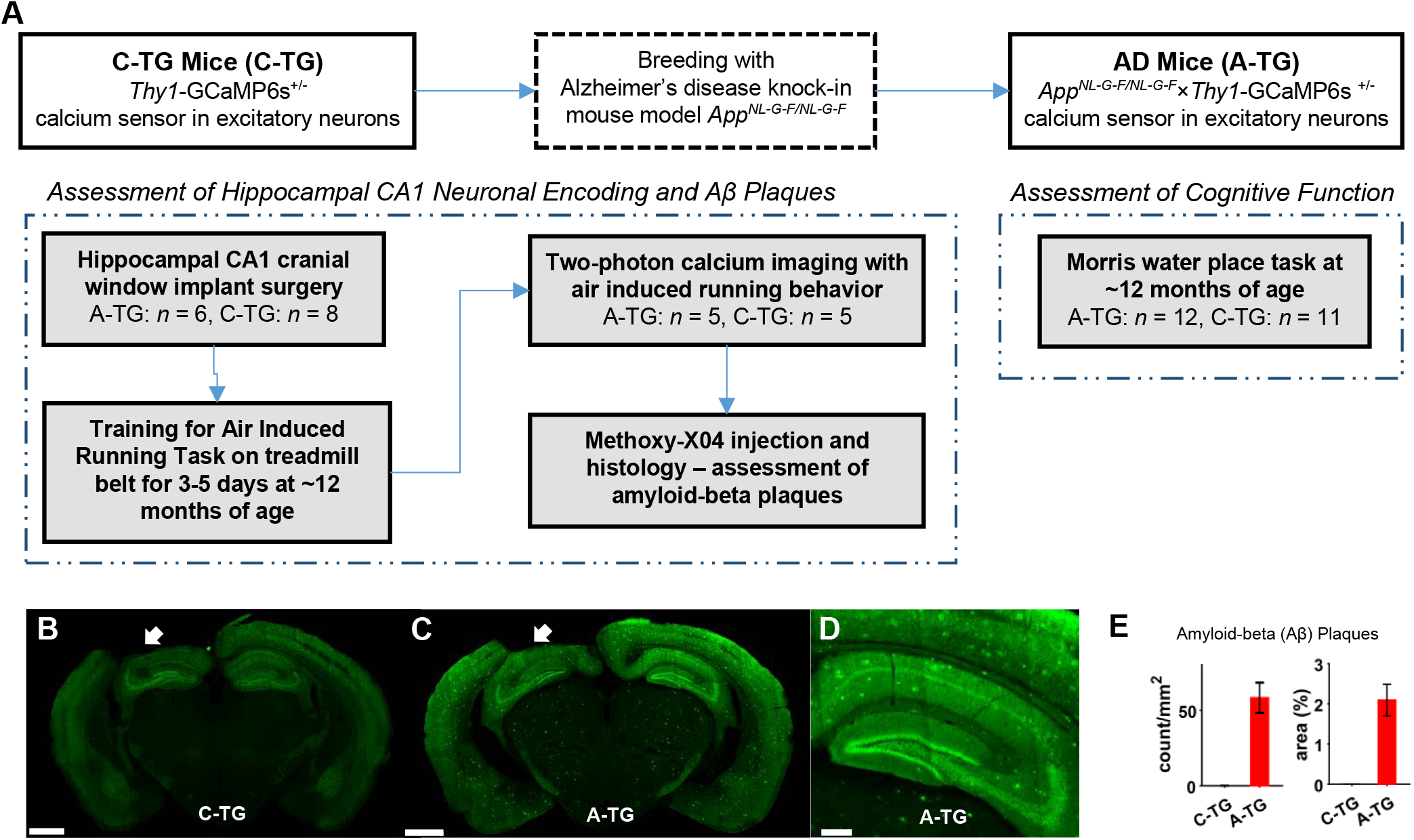
Experimental paradigm to test the role of familial Alzheimer disease (AD)-linked *APP* mutations on neuronal encoding and cognitive function in a knock-in mouse model. A) Generation of *App^NL-G-F/NL-G-F^×Thy1*-GCaMP6s (A-TG) mice to allow calcium imaging of neuronal activity, and experimental flow chart. B) Section from a 14-month-old C-TG mouse brain showing a lack of methoxy-X04 staining. The white arrow indicates the section of cortex that was removed by aspiration to allow imaging of CA1. Scale bar = 1 mm. C) Section from an 11-month-old A-TG mouse brain showing abundant labeling of amyloid-beta (Aβ) plaques with methoxy-X04. Labeling of plaques was clear, despite a high background level of green fluorescence in the hippocampus and neocortex due to GCaMP6 expression in neurons of these regions. Scale bar = 1 mm. D) High magnification view of the hippocampus from C. Scale bar = 250 μm. E) Quantification of Aβ plaque density (left) and percent area (right) in the hippocampus of C-TG and A-TG mice. C-TG: *n* = 6 mice; A-TG: *n* = 6 mice. Error bars indicate SEM.

## RESULTS

### A-TG and C-TG Mice Show Similar Behavioral Performance in an Air Induced Running Task

To study neuronal encoding of event-related distance and time in hippocampal CA1 neurons, we used an air induced running task for head-fixed mice (Inayat et al., 2022). The mice (8 C-TG, 6 A-TG) were trained to run a fixed distance on a conveyor belt in response to a stream of pressurized air delivered to the animal’s back (Fig. 2A). During training, the belt was devoid of cues, with the possible exception of olfactory cues deposited by the mouse during its training session. All mice were trained for 4-6 days to reach criterion (C-TG: average = 4.25 days; A-TG: average = 3.67 days). A trial consisted of an air-on phase and a 15 s air-off phase (Fig. 2B). Average speed during both phases was analyzed across the first 3 training days [three-way ANOVA, with trial phase (air-on vs. air-off) and day as repeated-measures factors] (Fig. 2C). There was no difference between the C-TG and A-TG mice (main effect of group; group × trial phase interaction; group × day interaction; three-way interaction: *p* > .25 for all). There was a significant interaction between training day and trial phase [*F*(2,24) = 20.32, *p* < .001]. Simple-effects tests showed that average speed during air-on increased significantly across days [*F*(2,11) = 19.84, *p* < .001], but speed during the air-off did not change [*F*(2,11) = 1.90, *p* = .561]. Follow-up pairwise comparisons showed that average speed during the trials increased significantly from day 1 to day 2 (*p* < .001) and from day 2 to day 3 (*p* = .049). In addition, speed was significantly greater during air-on than during air-off for all 3 days (*p* < .001 for each comparison). These results indicate that the mice learned to run in response to the air and learned not to run when it was off.

**Figure 2.**
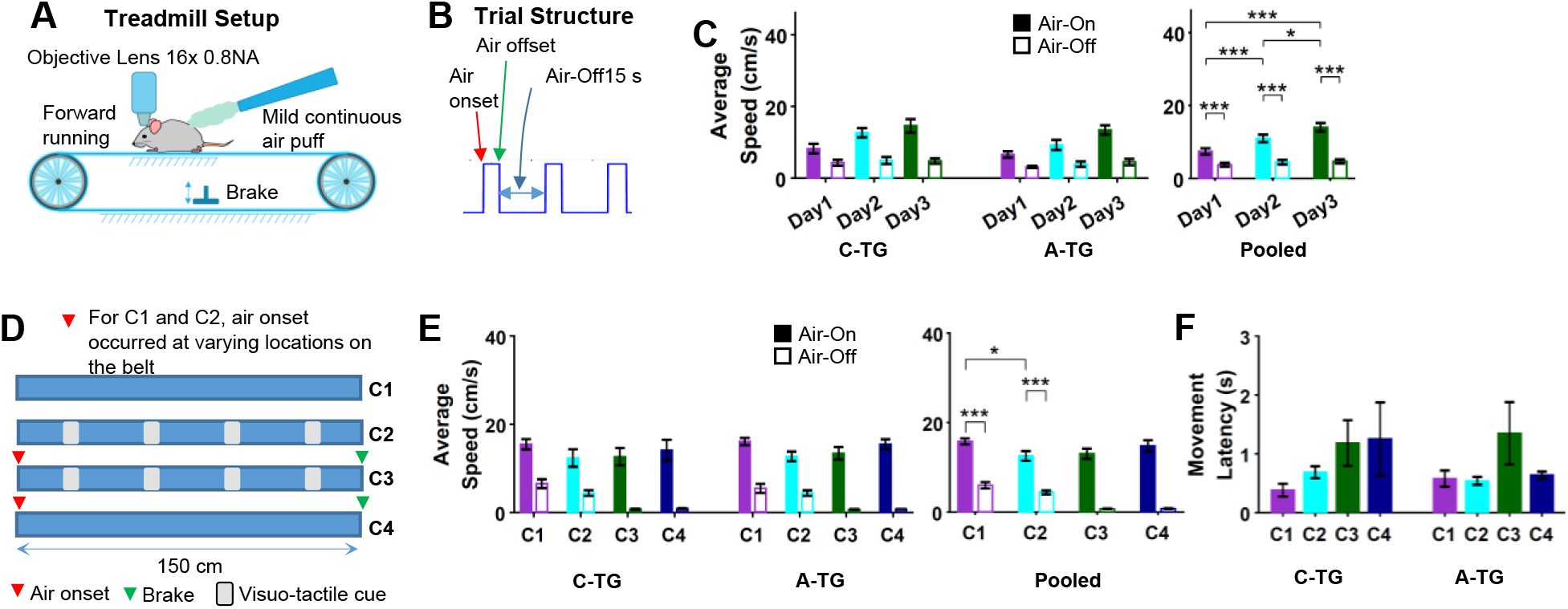
Behavioral performance of A-TG mice in the air induced running task was comparable to C-TG mice. A) Diagram of the non-motorized treadmill setup. B) Schematic of the trial structure. To initiate a trial, the air stream turned on and remained on until the mouse ran a fixed distance. After a 15 s interval with air-off, the air would come on again, initiating the next trial. C) Average running speed across training days for C-TG (*n* = 8) and A-TG (*n* = 6) mice [mixed ANOVA for examining the effect of Group, Day, and Trial Phase (Air-On vs. Air-Off)]. D) Schematic of the four belt conditions used for testing. Mice performed 10 trials with each belt configuration. Configuration 1 (C1) consisted of a blank belt with no cues. For Configuration 2 (C2) visuo-tactile cues were added to the belt. For Configuration 3 (C3) a brake was applied during the inter-trials, locking the cue locations to a specific distance relative to the air onset. Configuration 4 (C4) was the same as C1, but with the addition of the brake. E) Average running speed across the four belt configurations during testing/imaging (mixed ANOVA for examining the effect of Group, Configuration, and Trial Phase). F) Average latency to begin moving following air onset during testing/imaging (mixed ANOVA for examining the effect of Group and Configuration). For E and F, C-TG: *n* = 5 mice; A-TG: *n* = 5 mice. Error bars indicate SEM. \**p* < .05, \*\*\**p* < .001

Following training, CA1 neurons were imaged while the mice were given four belt configurations, two in which a mouse was free to walk on the belt during the air-off phase and two in which the break was applied to prevent walking during the air-off phase (Fig. 2D). Mice completed 10 trials of a configuration before moving onto the next. In Configuration 1, air-on served as the cue, and self-motion information provided the only source of information to track distance travelled. In Configuration 2, visuo-tactile cues were attached to the same belt, creating a richer sensory environment that changed as the belt moved. In Configuration 3, the brake was applied during the air-off intervals, preventing the mice from advancing the belt; thus, belt cues were consistently aligned with the air frame of reference. In Configuration 4, the brake was applied during the air-off intervals, and there were no cues (i.e., cues were removed from the belt).

To compare the performance of C-TG and A-TG mice, average movement speed was analyzed across the four belt conditions [three-way ANOVA, with trial phase (air-on vs. air-off) and configuration as repeated-measures factors]. There was no difference in speed between groups (main effect of group; group × trial phase interaction; group × configuration interaction; three-way interaction: *p* > .55 for all). There was a significant interaction between configuration and trial phase [*F*(3,24) = 16.33, *p* = .001] (Fig. 2E). Follow-up comparisons (Bonferroni-corrected paired *t*-tests of difference scores) showed that the difference in speed between trial phases was greater for Configuration 4 compared to Configuration 1 (*p* = .033), and greater for both Configuration 3 and 4 compared to Configuration 2 (*p* = .002, *p* < .001, respectively). This was not surprising, considering that speed was clamped at zero, by the brake, during the air-off intervals on Configurations 3 and 4, but mice were free to move during air-off intervals in Configurations 1 and 2. Despite this, speed was still significantly greater during the air-on phase than the air-off phase for both Configurations 1 and 2 (*p* < .001 for each comparison). Comparing average speed during the air-on periods across configurations, there was also a significant difference, with speed being greater on Configuration 1 than Configuration 2 (*p* = .024). This may be attributed to the addition of the visuo-tactile cues on the belt for Configuration 2, near which the mice slowed down to investigate, especially on the initial trials after the cues were added.

The latency to start running following the air onset (Fig. 2F) did not differ between the C-TG and A-TG mice [main effect of group: *F*(1,8) = 0.11, *p* = .748; group × configuration interaction: *F*(3,24) = 0.84, *p* = .443] or across conditions [*F*(3,24) = 2.93, *p* = .089]. Though the effect was not significant, the mice did tend to take longer to start running on the Configuration 3 trials compared to Configurations 1 and 2. This was likely because, as the mice were prevented from running during the air-off periods, they may have taken longer to initiate running as the brake was released.

### Hippocampal CA1 Neurons in A-TG mice Exhibit Diminished Average Firing Rates

There was no significant difference between C-TG and A-TG mice in the total number of CA1 neurons that were detected by calcium imaging (Fig. 3A-C). Nevertheless, when we examined the firing rate of each neuron, inferred from the calcium trace by deconvolution (Fig. 3D), the average firing rate of the cells differed between the C-TG and A-TG mice across the entirety of the recording. The average firing rate in A-TG mice was significantly lower compared to C-TG mice [*t*(8) = 2.63, *p* = .030] (Fig. 3E). When we partitioned the recordings based on whether the mouse was at rest (speed = 0) or in motion (speed > 0) to assess the influence of movement on firing rate [two-way ANOVA, with behavioral state (motion vs. rest) as a repeated-measures factor] (Fig. 3F), firing rate increased significantly with motion [*F*(1,8) = 58.51,*p* < .001]. There was, however, no significant difference in firing rate between the C-TG and A-TG mice [main effect of group: *F*(1,8) = 4.34, *p* = .071; group × behavioral state interaction: *F*(1,8) = 5.10, *p* = .054]. We also analyzed the relative change in firing rate between motion and rest (by calculating the percent increase). There was no significant difference between the C-TG and A-TG mice [*t*(8) = −0.11, *p* = .912]. Firing rate increased in both groups by about 37% when the mice were moving compared to when they were at rest (C-TG: 36.5 ± 7.0%; A-TG: 37.5 ± 5.5%). However, when the firing rate was compared between the two groups across test configurations and phases of the air application of air (Air-On vs. Air-Off) with a three-way ANOVA, significant main effects of Group and Air-Phase were observed (Fig. 3G) [main effect of group: *F*(1,8) = 7.26, *p* = .027, η^2^ = .40, main effect of Air-Phase: *F*(1,1) = 31.14,*p* < .001, η^2^ = .15]. Overall, these results suggest that A-TG mice had hypoactive neurons irrespective of the behavioral state (rest or motion) or configuration (C1-C4).

**Figure 3.**
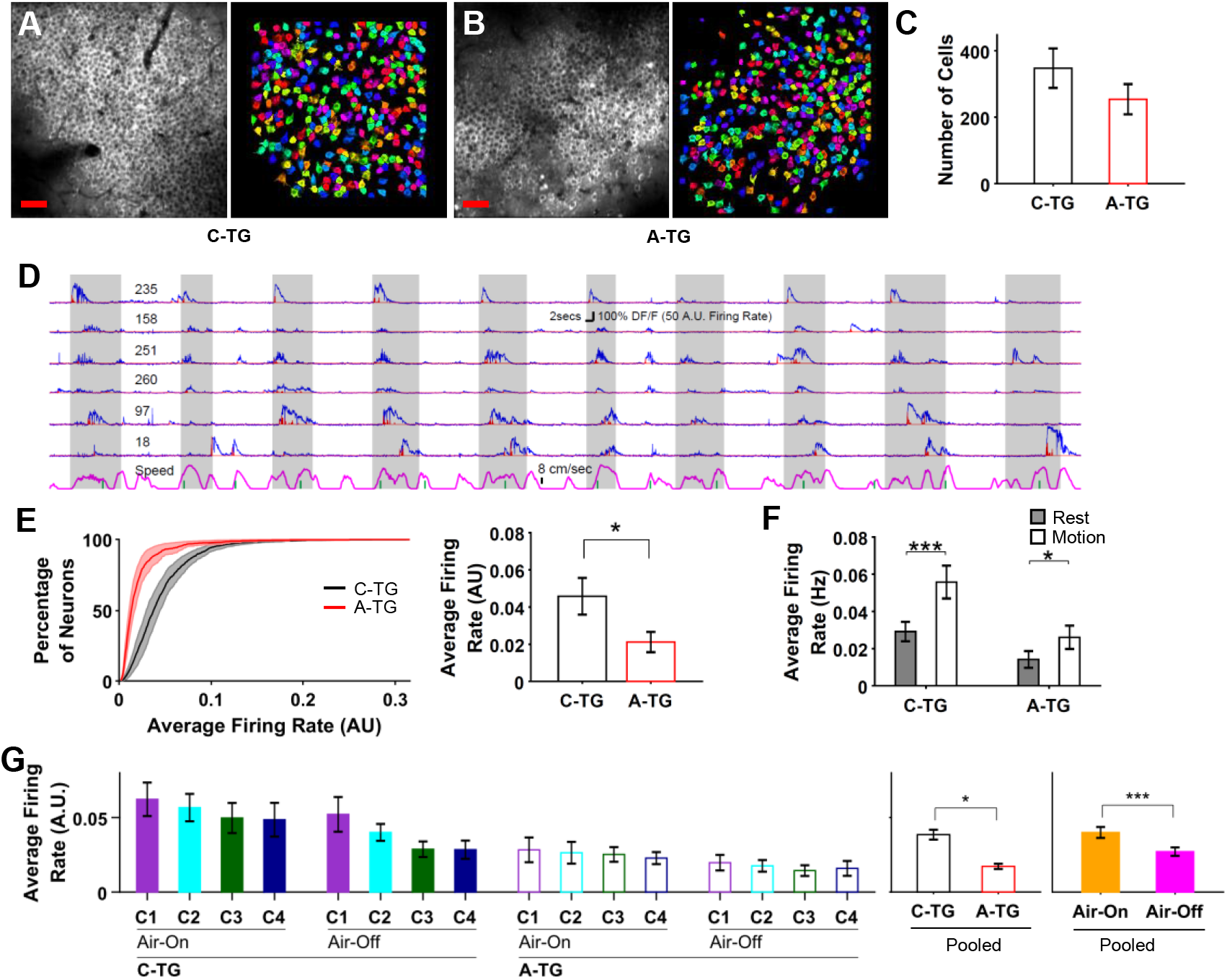
Calcium imaging of hippocampal CA1 neurons revealed decreased firing rate in A-TG compared to C-TG mice. A, B) Average image of a calcium imaging stack for C-TG and A-TG mice, with randomly color-coded regions showing identified neurons. Scale bar = 50 μm. C) The mean number of neurons detected in the C-TG and A-TG mice did not differ. D) ΔF/Fo (blue traces) and deconvolved firing rate (red traces) of representative neurons from one C-TG mouse are shown, with gray regions indicating application of the air stream. Magenta line at the bottom indicates mouse speed. E) Left: Cumulative distributions, averaged across mice, of the firing rate (averaged across the entire recording) of CA1 neurons in C-TG and A-TG mice. Right: the average firing rate of CA1 neurons in A-TG mice was significantly lower compared to C-TG mice. F) Average firing rate of CA1 neurons when the mice were at rest (running speed = 0) or in motion (running speed > 0). In both groups, firing rate increased by about 37% when the mice were in motion. In both groups, firing rate increased by about 37% when the mice were in motion. G) Average firing rate of CA1 neurons in Configurations C1-C4 and during Air-On and Air-Off periods for both C-TG and A-TG mice (mixed ANOVA with Groups, Configurations, and Air-Phase as factors). C-TG: *n* = 5; A-TG: *n* = 5. Error bars indicate SEM. \**p* < .05, \*\*\**p* < .001

### Responsive Cells in A-TG and C-TG Mice Express Similar Distance and Time Tuning Characteristics

We compared the spatial and temporal tuning characteristics of cells in A-TG and C-TG mice by identifying cells whose activity was modulated by traveled distance from air onset or elapsed time from air offset.

#### Distance-tuned cells

For identifying distance tuning from air onset, peri-event distance raster plots (PEDRs) and histograms (PEDHs) were generated for each neuron (Fig. 4A). A neuron was deemed as responsive if there was a significant change in the mean firing rate across raster bins (distance) detected using a set of independent one-way ANOVAs (see methods). To qualitatively assess the responses of all responsive cells, rate vector maps (RVMs, top rows in Fig. 4C and 4D) showing normalized PEDHs of all responsive cells were generated. RVMs showed transient firing of cells over the length of the belt and were not qualitatively different for C-TG and A-TG mice. This was confirmed by plotting and observing population vector correlation plots (middle and bottom rows, Fig. 4C and 4D) in which each pixel shows a Pearson correlation between two population vectors (i.e., two columns of a RVM). The correlation plots showed elevated densities concentrated around the diagonal that were similar for both C-TG and A-TG mice, indicating transient firing of cells along the length of the belt.

**Figure 4.**
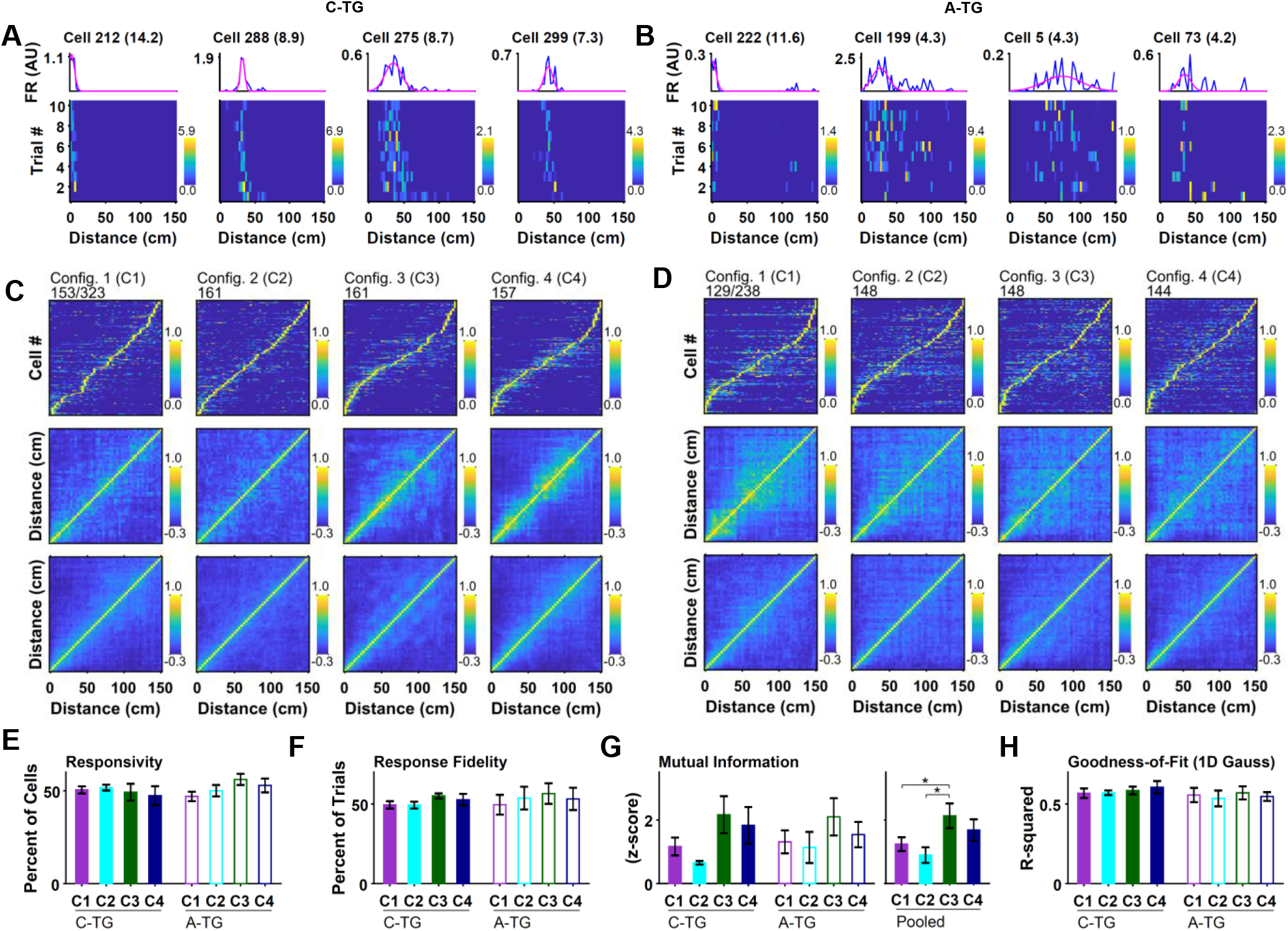
Event-related distance encoding of hippocampal CA1 neurons was similar in A-TG and C-TG mice. A) Raster plots showing firing rate based on trial number and distance traveled from the air stream onset (bottom row) and peri-event distance histograms (top row) for selected cells from C-TG mice. B) Same as A but for A-TG mice. C) Rate vector maps (top row) showing neuronal response as a function of distance travelled from the air stream onset, averaged across trials (normalized by peak response), for all responsive neurons from a representative C-TG mouse for all four configurations, C1-C4. Population vector correlation plots (middle row), in which each pixel represents the Pearson correlation coefficient of two columns (bins) in the rate vector map. The glow around the diagonal indicates spatially-localized transient firing of neurons. Average population vector correlation plots (bottom row) for the entire group of 5 mice. D) Same as C but for A-TG mice. E-H) Responsivity, response fidelity, mutual information (z-scored), and goodness-of-fit of a 1D Gaussian, respectively, calculated from the population of responsive neurons for C-TG and A-TG mice across the four configurations (mixed ANOVA with Group and Configuration as factors). Error bars indicate SEM. *\*p* < .05

Next, the percentage of responsive cells (responsivity, Resp) and their properties, including response fidelity (RF), mutual information (MI), and goodness-of-fit (GOF) of a 1D Gaussian, were quantitatively compared between A-TG and C-TG mice. RF was defined as the percentage of trials in a PEDR in which the cell’s firing rate was greater than zero at any point during a trial. MI between firing rate and distance was determined from a PEDR (Souza et al., 2018) and then z-scored (zMI) using a distribution of 500 MI values calculated from randomly shuffled instances of the same PEDR. A larger zMI value mostly indicates better spatial repeatability of a cell’s responses across trials (i.e., highly tuned cells). GOF was measured as the R-squared value for a 1D Gaussian least-squares fit on the PEDH (magenta lines in Fig. 4A). Whereas the first two measures quantified the robustness of responses across trials, the third measure assessed whether the firing rate transiently increased and decreased with respect to distance in a Gaussian manner, assuming a single tuning profile (observed for most of the cells from the population vector plots).

The Resp, RF, and GOF compared between the C-TG and A-TG groups (by setting Configuration as a within-subjects factor) was not different, and neither was the Group × Configuration interaction (all p values > 0.05, Fig. 4E, F, and H). However, there was a significant effect of configuration on zMI [*F*(3,24) = 10.32, *p* < 0.001, η^2^ = .21], with *post hoc* tests revealing a larger mean zMI for Configuration 3 compared to 1 and 2 (Fig. 4G). This was because of the more consistent trial-to-trial environment presented to mice in Configuration 3.

From the set of responsive cells found earlier, further analysis was done on highly distance-tuned cells (with zMI > 1.65) to examine properties of cells active at different distances from air onset (Fig. 5A and 5B). For this analysis, distance was binned into three bins, B1, B2, and B3, corresponding to distances of 0-50, 50-100, and 100-150 cm. Each responsive cell was placed into one of the bins based on the location of the center of the fitted 1D Gaussian. The Resp, RF, zMI, GOF, and tuning field width (FW) were then compared between A-TG and C-TG mice by setting Bin and Configuration as within-subjects factors. No Group effect was observed for any of these factors. However, for responsivity, there were main effects for both Bin [*F*(2,2) = 24.45,*p* < .001, η^2^ = .56] and Configuration [*F*(3,3) = 9.96, *p* = .003, η^2^ = .14]. *Post hoc* tests revealed a larger percentage of cells in B1 compared to B2 and B3, and in C3 compared to C2 and C4 (Fig. 5C). For RF, there was a main effect of Bin [*F*(2,2) = 6.30, *p* = .022, η^2^ = .12] but no significant *p-*values from the *post hoc* tests (Fig. 5D). For zMI, there were main effects of Bin [*F*(2,2) = 6.00, *p* = .023, η^2^ = .24] and Configuration [*F*(3,3) = 8.86, *p* = .003, η^2^ = .14] (Fig. 6E). For GOF, there was a main effect of Bin [*F*(2,2) = 15.34,*p* < 0.001, η^2^ = .32], and for FW there was a main effect of Configuration [*F*(3,3) = 5.94, *p* = .012, η^2^ = .05] (Fig. 5F and 5G). Overall, the results from highly tuned cells were similar to those obtained by considering all distance-tuned cells and no group differences were found.

**Figure 5.**
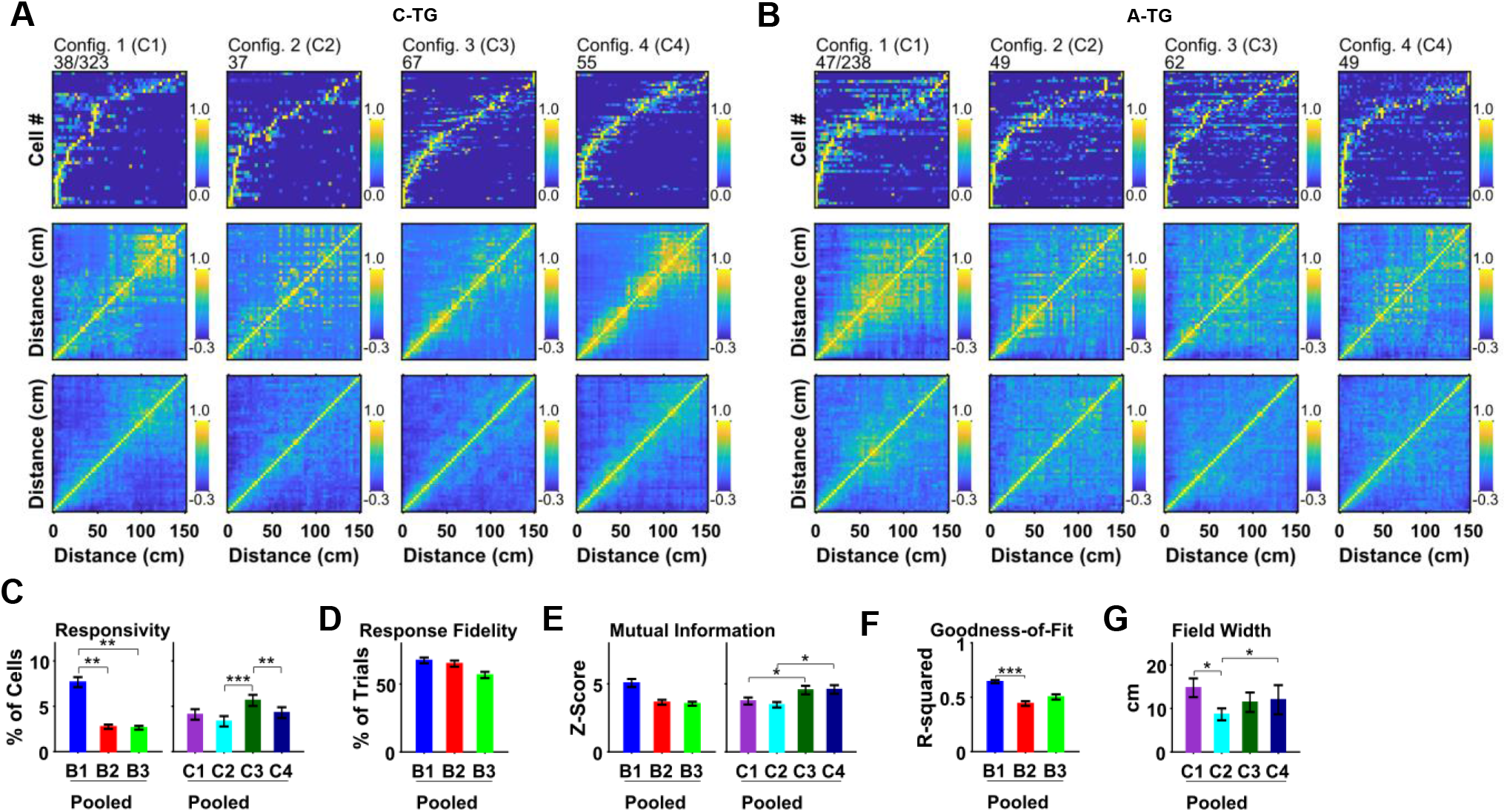
The distance-encoding properties of highly tuned (zMI > 1.65) hippocampal CA1 neurons were similar in C-TG and A-TG mice. A) Rate vector maps (top row) showing neuronal response as a function of distance travelled from the air stream onset, averaged across trials (normalized by peak response), for highly tuned responsive neurons from a representative C-TG mouse for all four configurations, C1-C4. Population vector correlation plots (middle row), in which each pixel represents the Pearson correlation coefficient of two columns (bins) in the rate vector map. The glow around the diagonal indicates spatially-localized transient firing of neurons. Average population vector correlation plots (bottom row) for the entire group of 5 mice. B) Same as A but for A-TG mice. C-G) Responsivity, response fidelity, mutual information (z-scored), and goodness-of-fit of a 1D Gaussian, respectively, calculated from the population of highly tuned responsive neurons for C-TG and A-TG mice across the four configurations (mixed ANOVA with Group and Configuration as factors). Error bars indicate SEM. \**p* < .05, \*\**p* < .01, \*\*\**p* < .001

**Figure 6.**
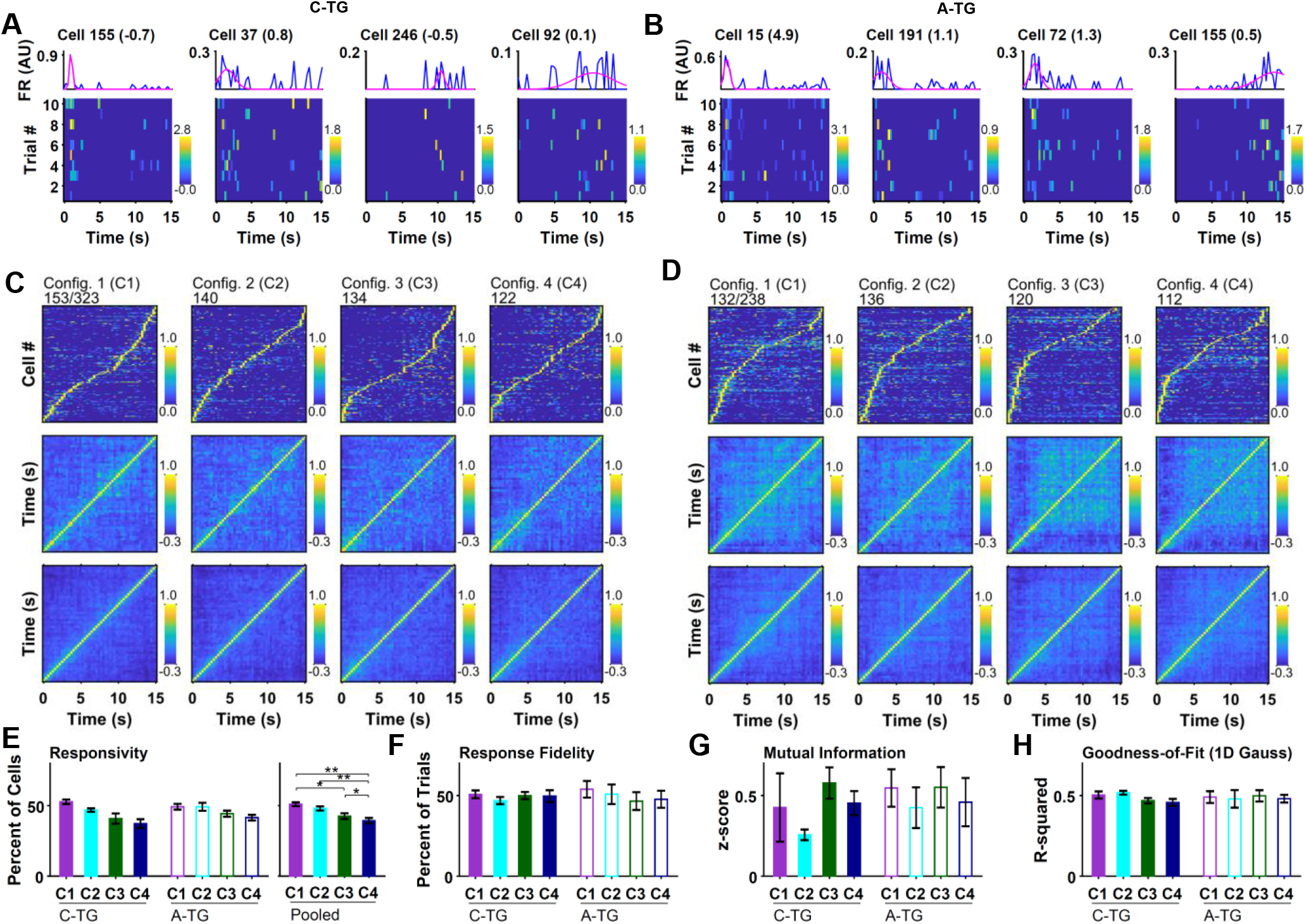
Event-related time encoding of hippocampal CA1 neurons was similar in A-TG and C-TG mice. A) Raster plots showing firing rate based on trial number and time from the air stream offset (bottom row) and peri-event time histograms (top row) for selected cells from C-TG mice. B) Same as A but for A-TG mice. C) Rate vector maps (top row) showing neuronal response as a function of time from the air stream offset, averaged across trials (normalized by peak response), for all responsive neurons from a representative C-TG mouse for all four configurations, C1-C4. Population vector correlation plots (middle row), in which each pixel represents the Pearson correlation coefficient of two columns (bins) in the rate vector map. The glow around the diagonal indicates temporally-localized transient firing of neurons. Average population vector correlation plots (bottom row) for the entire group of 5 mice. D) Same as C but for A-TG mice. E-H) Responsivity, response fidelity, mutual information (z-scored), and goodness-of-fit of a 1D Gaussian, respectively, calculated from the population of responsive neurons for C-TG and A-TG mice across the four configurations (mixed ANOVA with Group and Configuration as factors). Error bars indicate SEM. \**p* < .05, \*\**p* < .01

#### Time-tuned cells

Time-tuned cells from air offset during air-off intervals were next analyzed, in a similar manner as was done for distance-tuned cells. Peri-event time raster plots (PETRs) and histograms (PETHs) were generated with air offset as the reference point. Raster plots (Fig. 6A and 6B), RVMs, and population vector correlations (Fig. 6C and 6D) showed no qualitative difference between the C-TG and A-TG groups. Quantitatively, RF, zMI, and GOF compared between the C-TG and A-TG groups were not significantly different (Fig. 6F-H). However, the Resp was different across configurations [*F*(3,24) = 17.01, *p* < 0.001, η^2^ = .47], with a smaller percentage of cells being active in each successive configuration (Fig. 6E).

Next, the properties of highly tuned cells (zMI > 1.65) were compared between A-TG and C-TG mice by binning the air-off duration of 15 s into three bins, B1, B2, and B3, corresponding to 0-5, 5-10, and 10-15 s. Cells were placed in one of the three bins based on the center of the fitted Gaussian. The Resp, RF, zMI, GOF, and FW were then compared for the two Groups, with Bin and Configuration considered as within-subjects factors. There was no main effect of Group for any of the variables except for GOF, for which there was a Group × Bin × Configuration interaction [*F*(6,36) = 3.72, *p* = .032, η^2^ = .19]. For Resp, there were main effects of both Bin [*F*(2,2) = 7.44, *p* = .016, η^2^ = .18] and Configuration [*F*(3,3) = 4.53, *p* = .021, η^2^ = .07] (Fig. 7C). For RF, there was significant Bin × Configuration interaction [*F*(6,6) = 3.79, *p* = .029, η^2^ = .09], and for zMI, there was main effect of Bin [*F*(2,2) = 4.89, *p* = .044, η^2^ = .09] (Fig. 7D and 7E). There was no significant effect of any factor nor a significant interaction for FW. These results show that the responsive cells, including cells with robust and highly tuned responses, have similar properties in the A-TG and C-TG mice.

**Figure 7.**
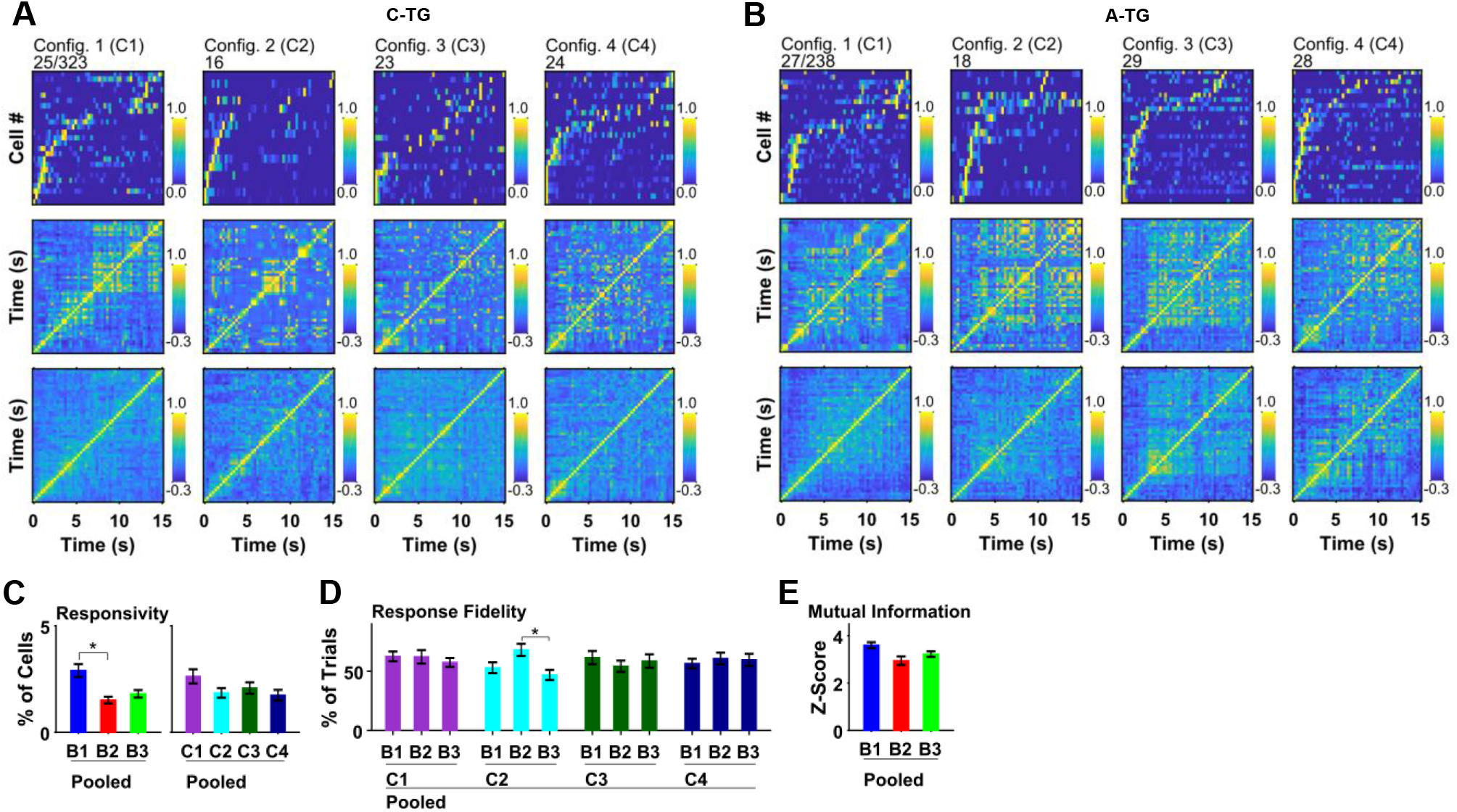
The time-encoding properties of highly tuned (zMI > 1.65) hippocampal CA1 neurons were similar in C-TG and A-TG mice. A) Rate vector maps (top row) showing neuronal response as a function of time from the air stream offset, averaged across trials (normalized by peak response), for highly tuned responsive neurons from a representative C-TG mouse for all four configurations, C1-C4. Population vector correlation plots (middle row), in which each pixel represents the Pearson correlation coefficient of two columns (bins) in the rate vector map. The glow around the diagonal indicates temporally-localized transient firing of neurons. Average population vector correlation plots (bottom row) from the entire group of 5 mice. B) Same as A but for A-TG mice. C-E) Responsivity, response fidelity, mutual information (z-scored), and goodness-of-fit of a 1D Gaussian, respectively, calculated from the population of highly tuned responsive neurons for C-TG and A-TG mice across the four configurations (mixed ANOVA with Group and Configuration as factors). Error bars indicate SEM. \**p* < .05

### The Remapping Dynamics of Responsive Cells Across Test Configurations are not Different Between A-TG and C-TG Mice

The changes in cellular activity in response to transitions between pairs of behavioral configurations were analyzed and compared between A-TG and C-TG mice, as an assessment of short-term cellular plasticity pertaining to the distance to be run or time to wait for the next air trial. All responsive cells in either of the configurations were included in this analysis. Three parameters, including the activity correlation (spatial or temporal), the population vector (PV) correlation, and the change in firing rate (rate remapping), were calculated (see methods). The activity correlation quantified the similarity of responses of cells across configurations (i.e., whether the firing activity of a cell was similar at the same distance or time from configuration to configuration). The population vector correlation quantified the similarity of responses of the whole population (all cells) bin wise (i.e., whether the activity of all cells in a bin was similar across configurations). The rate remapping score quantified, for each cell, the changes in firing rate across bins from configuration to configuration (i.e., whether the tuning of a cell remained the same across configurations but only firing rate changed). The three parameters were calculated for three transitions: from Configuration 1 to Configuration 2 (C12), Configuration 2 to Configuration 3 (C23), and Configuration 3 to Configuration 4 (C34). The transition was set as a within-subjects factor in the ANOVA. This analysis was done for all responsive cells, as well as separately for highly tuned cells.

For distance-tuned cells, there was no main Group effect or Group × Transition interaction for any of the dependent variables, regardless of whether all responsive cells or only highly tuned cells were considered (Fig. 8A-E). There was a main effect of Transition (see figure), with *post hoc* tests revealing that cellular responses were more stable for the C34 transition compared to C23 or C12. For time-tuned cells, there was no significant Group effect or Group × Transition interaction for any of the variables, except for a Group × Transition interaction for the PV correlation when all responsive cells were considered (Fig. 8G). The main effect of Transition showed a higher temporal correlation for the C12 and C34 transitions compared to C23 (Fig. 8F and 8I), and a higher rate remapping score for C23 compared to C12 and C34, suggesting robust cellular activity transitions for the C12 and C34 transitions. These results show that the dynamics of cellular activity across configurations were not different between A-TG and C-TG mice. Consistent with the nature of the behavioral task, however, transition C34 showed relatively stable activity of cells (less remapping) even when the visuo-tactile cues were removed, showing robust/stable short-term cellular plasticity.

**Figure 8.**
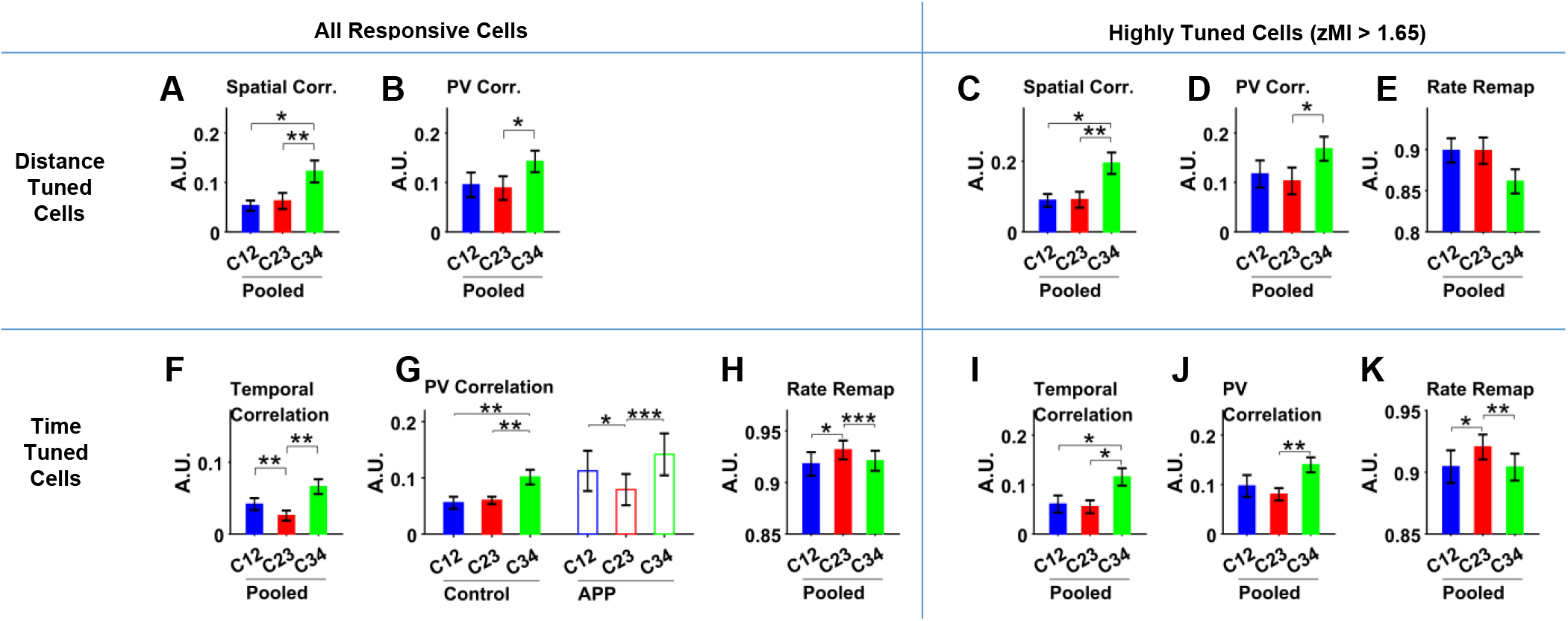
Remapping of CA1 neuronal populations encoding event-related distance and time was similar in C-TG and A-TG mice. Remapping scores are shown for transitions between contiguous configurations (C12 indicates the transition from Configuration 1 to Configuration 2, etc.). Except for in G, only results pooled across Group (C-TG and A-TG) are shown, because there was no significant effect of Group or Group × Transition interaction on remapping. A-K) Spatial/temporal correlation (Corr.) scores, population vector correlation scores, and rate-remapping scores for both distance- (A-E) and time- (F-K) tuned responsive CA1 neurons for C-TG and A-TG mice across the four configurations (mixed ANOVA with Group and Transition as factors). No significant effect was seen for rate-remapping scores for distance-tuned cells. Error bars indicate SEM. \**p* < .05, \*\**p* < .01, \*\*\**p* < .001

### Similar Dynamics in the Recruitment of CA1 Neurons Across Test Configurations in A-TG and C-TG Mice

The recruitment of cells across changing behavioral configurations was analyzed by finding the conjunction and complementation of cellular populations that were deemed responsive in individual configurations. The conjunction of populations in two configurations identified cells that were active in both configurations (C **∈** A and B), whereas the complementation of two populations identified cells that were not shared (C1 **∈** A not B or C2 **∈** B not A). The conjunction and complementation of all pairs of responsive populations of distance- and time-encoding cells in the 4 configurations were considered (Fig. 9A and 9B). For both A-TG and C-TG mice, the maximum conjunction of cells was between distance-tuned cells in Configurations 3 and 4. Overall, the conjunction was greater between similarly tuned cells. For instance, the conjunction was greater between distance-tuned cells compared to between distance- and time-tuned cells. Conversely, the complementation of distance- and time-tuned cells was greater compared to complementation between similarly tuned (distance or time) cells. An agglomerative hierarchical clustering of all the conjunction and complementation values was done to identify similarity of Configurations/Tuning-Type (Fig. 9C and 9D) e.g., two configurations would be similar to each other if their conjunction values with all other configurations are similar (see methods). For both groups, configurations were clustered separately by Tuning-Type (distance or time; separation of red and cyan clusters in Fig. 9C and 9D). Furthermore, C1-D was closer to C2-D, whereas C3-D was closer to C4-D, corresponding to the No-Brake and Brake conditions, respectively. The clustering of time-tuned cells was also similar for both groups. The clustering analysis produced even a clearer separation between configurations and tuning-type when only highly tuned cells were considered (Fig. 9E and 9F). These results suggest that the recruitment of distance and time tuned cells across configurations was similar between the A-TG and C-TG mice.

**Figure 9.**
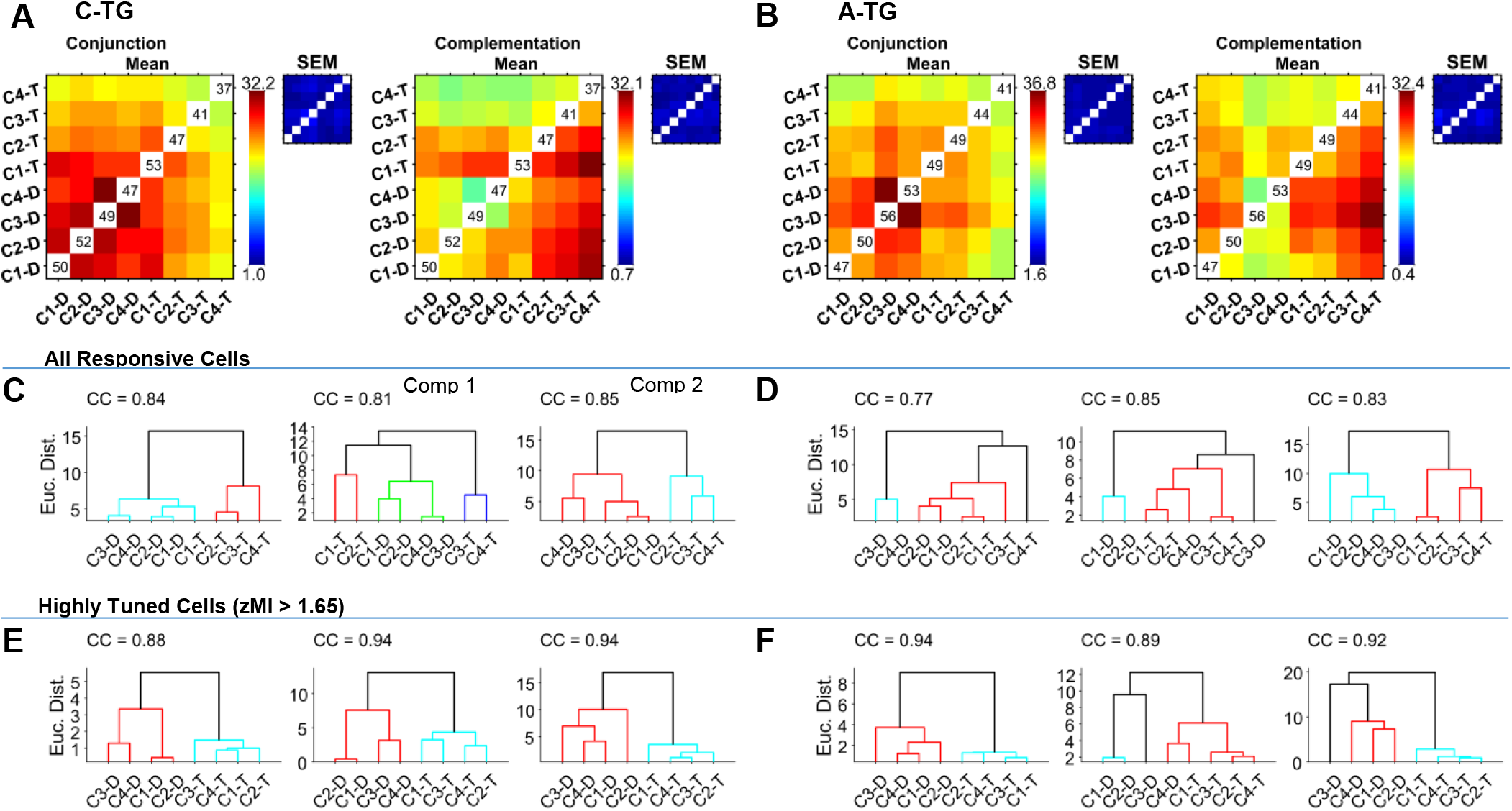
Conjunction and complementation of populations of responsive CA1 neurons across behavioral configurations was similar in A-TG and C-TG mice. A) Mean and SEM (insets) heatmaps (over 5 C-TG mice) of the percentages of conjunction and complementation for all pairs of responsive neuronal populations from the four belt configurations, for both the time (T) and distance (D) encoding populations (e.g., C1-T and C1-D indicate time and distance encoding, respectively, in Configuration 1). A higher percentage of conjunction between two populations indicates a larger overlap of responsive neurons. A higher percentage of complementation indicates a larger percentage of responsive neurons in an individual event. Complementation 1 between populations X and Y corresponds to neurons active in X but not Y whereas Complementation 2 corresponds to neurons active in Y but not in X. (B) Same as A but for A-TG mice. C) Dendrograms resulting from agglomerative hierarchical clustering of the average conjunction (left) and complementation (middle and right corresponding to complementation 1 and 2) heatmaps/matrices of C-TG mice, obtained for all responsive cells. D) Same as C but for A-TG mice. E, F) Same as C and D, respectively, but when only highly tuned cells were considered. CC: cophenetic correlation as a measure of the reliability of dendrogram construction.

### Distance and Time are Decoded with Similar Accuracy in A-TG and C-TG Mice

The encoding of distance from air onset by the sampled population of cells in each animal was compared between A-TG and C-TG mice. All identified cells were considered for this analysis. Decoding was conducted by maximum a posteriori estimation of distance from cellular activity, while the accuracy of decoding was assessed by leave-one-out cross-validation (see methods; Fig. 10A and 10B). The average decoding error over distance is shown for A-TG and C-TG mice in Fig. 10C, illustrating similar trends for all configurations. There was no significant Group effect, but the decoding error was significantly different across configurations [*F*(3,3) = 6.19, *p* = .003, η^2^ = .19], with *post hoc* comparisons revealing a significantly larger error for C2 compared to C1 and C3 (Fig. 10D). The dynamics of encoding of distance by the neuronal population were also tested by training the model with activity in one configuration and testing with the activity of cells in the immediately subsequent configuration. There was no significant Group effect, but decoding error was significantly different across configuration transitions [*F*(2,2) = 7.14, *p* = .006, η^2^ = .24], with *post hoc* comparisons revealing a larger error for transition C12 compared to C23 and C34 (Fig. 10E).

**Figure 10.**
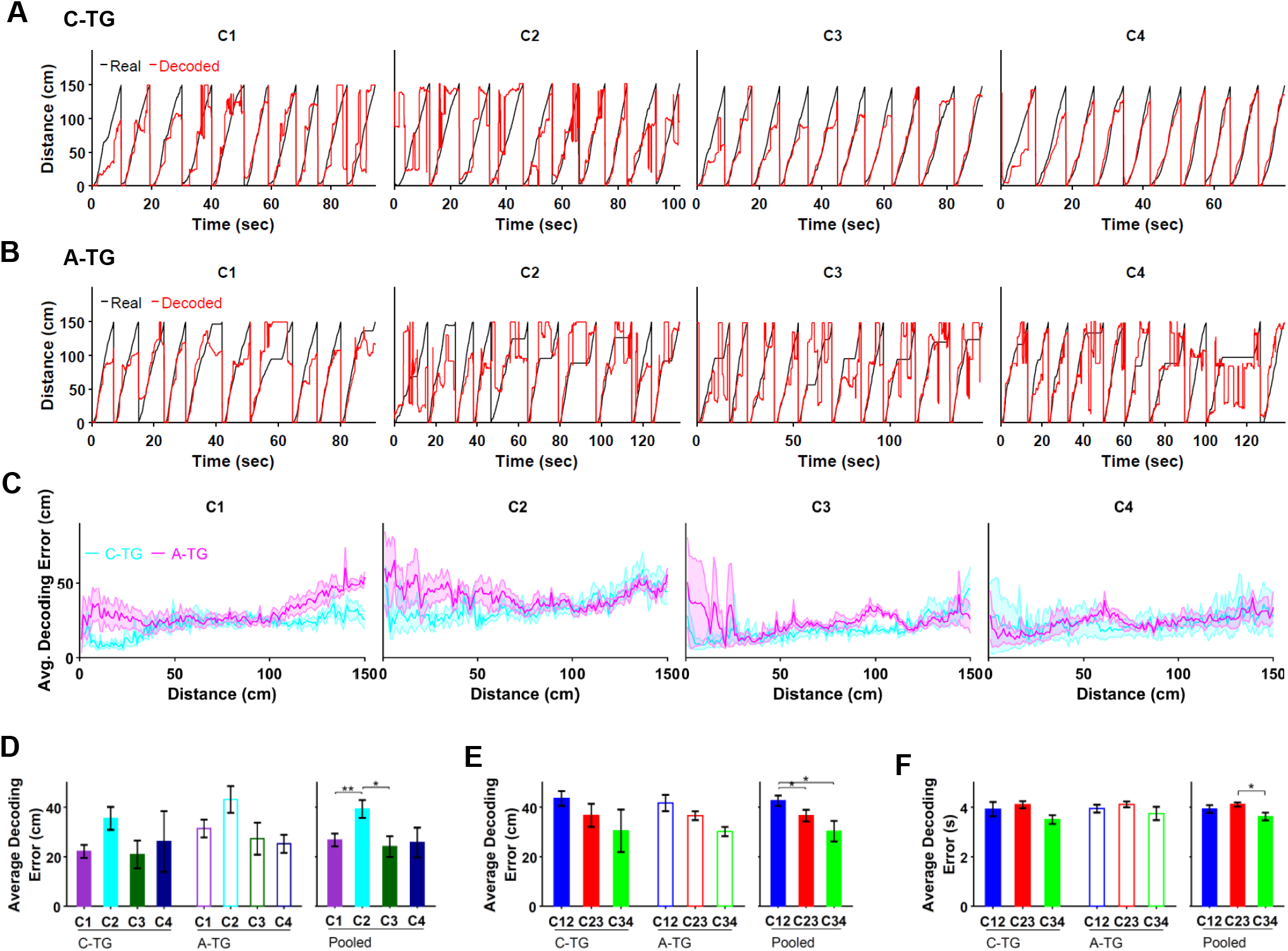
Bayesian decoding of distance and time from neuronal activity provides further evidence of similar distance and time encoding in A-TG and C-TG mice. All neurons were considered in this analysis. A) Real and decoded distance versus time (shown for all trials) in the four configurations, C1-C4, for a representative C-TG mouse. B) Same as A but for an A-TG mouse. C) Average decoding error versus distance traveled from the air stream onset. D) Average decoding error for C-TG and A-TG mice across the four configurations (mixed ANOVA with Group and Configuration as factors). E) Average decoding error when decoding for a given configuration was done using the neuronal activity from the previous configuration (mixed ANOVA with Group and Configuration Transition as factors). For example, C12 indicates that distance in C2 was decoded based on activity in C1 to measure the degree of remapping. F) Same as E but for time encoding. Error bars indicate SEM. \**p* < .05, \*\*\**p* < .001

The encoding of time from air offset by the sampled population of cells was compared between A-TG and C-TG mice. There was no significant effect of any factor (Group or Configuration) or Group × Configuration interaction on average decoding error. However, when the dynamics of encoding of time across configurations was considered, there was significant effect of Configuration Transition [*F*(2,2) = 5.30, *p* = .017, η^2^ = .20], with *post hoc* tests showing a larger decoding error for transition C23 compared to C34 (Fig. 10F). These results suggest that the neuronal encoding of distance and time in A-TG and C-TG mice is similar when the analysis was done by considering all the sampled neurons, in contrast to the aforementioned analyses, where a subset of neurons deemed as responsive (or responsive plus highly tuned) were considered.

### A-TG Mice Exhibit Impairment at Locating the Escape Platform in the Morris Water Place Task

The Morris water place task (MWT) was used to determine whether A-TG mice showed deficits similar to those reported in the original APPki strain (Mehla et al., 2019). The mice were trained for 8 days, with a probe test on the 9th day. The performances of approximately 12-month-old A-TG mice (*n* = 12, 6 males, mean age = 364 days, SD = 9 days, range = 356-378 days) and similarly aged controls (*n* = 11, 8 males, mean age = 392 days, SD = 32 days, range = 360-444 days) were compared with a two-way ANOVA, with day as a repeated-measures factor. Swim speed did not differ between the two groups [main effect of group: *F*(1,21) = 0.55, *p* = .468; day × group interaction: *F*(7,147) = 1.94, *p* = .106; Fig.11A]. Both C-TG and A-TG mice showed performance improving across days (Fig. 11B; C-TG: *p* < .001; A-TG: *p* < .001). Averaging across all days, time to find the platform was significantly longer for the A-TG mice than for the C-TG mice [main effect of group: *F*(1,21) = 12.72, *p* = .002], suggesting an impairment in spatial memory in the A-TG mice. There was no significant day × group interaction [*F*(7,147) = 0.88, *p* = .490].

**Figure 11.**
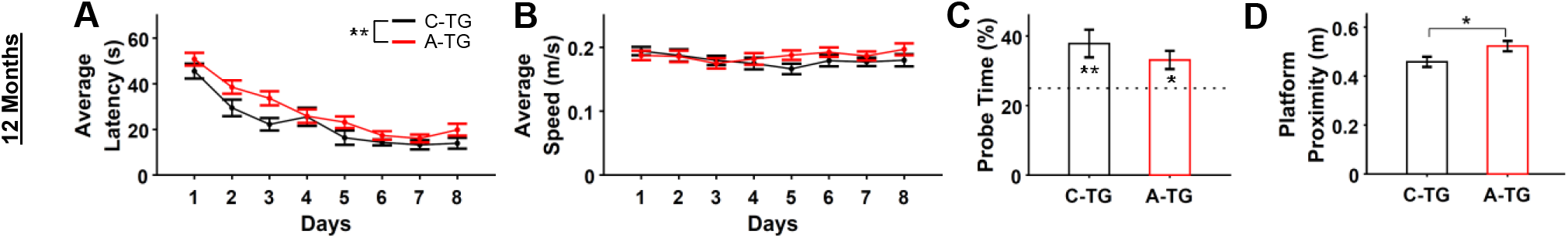
A-TG mice were mildly impaired at performing the Morris water place task (MWT) compared to C-TG mice. A) Averaging across all acquisition days, latency to find the platform was significantly greater in the A-TG mice than in the C-TG mice. B) Average swimming speed across the 8 days of acquisition training did not differ between the C-TG and A-TG mice. C) Memory for the platform’s location was assessed via a probe trial, conducted 24 h after the final acquisition day. There was no difference between the C-TG and A-TG mice in the percentage of time spent in the target quadrant, and both groups spent more time in the target quadrant than would be predicted by chance. D) During the probe trail, the average distance from the platform’s previous location was greater in the A-TG mice than the C-TG mice. Error bars indicate SEM. \**p* < .05, \*\**p* < .01

On the probe trial, there was no significant difference between the C-TG and A-TG mice when performance was assessed based on the percentage of total search time spent in the pool quadrant in which the platform was previously located [one-way ANOVA: *F*(1,21) = 1.03, *p* = .321; Fig. 11C]. One-sample *t*-tests confirmed that the percentage of time spent in the correct quadrant was significantly greater than would be predicted by chance (i.e., 25%) in both the C-TG mice [*t*(10) = 3.24, *p* = .009] and the A-TG mice [*t*(11) = 3.10, *p* = .010]. We next examined probe trial performance using a different measure: the average proximity to the platform’s previous location (Fig. 11D). In this case, there was a significant difference between groups, with the A-TG mice exhibiting a greater average distance from the target location compared to the C-TG mice [*F*(1,21) = 4.78, *p* = .040]. In summary, the probe test results supported our conclusions from the training trials. Both groups were able to learn and remember the platform location, but the A-TG mice exhibited a mild impairment in spatial memory.

## DISCUSSION

This study investigated cellular-level mechanisms that may contribute to the deficits in movement control and spatial navigation observed in AD. To enable the large-scale functional imaging of CA1 neuronal populations, A-TG mice were generated by crossing the APPki line with C-TG mice. At 12 months of age, A-TG mice had numerous Aβ plaques in the hippocampus (and throughout the brain, though, in this study, plaque load was quantified only in the hippocampus). The AT-G mice were not impaired in performing the head-fixed air-induced running task but their acquisition of a swimming pool place was impaired. The functional encoding of distance and time by CA1 neuronal populations was unimpaired in the A-TG group, as assessed through the air-induced running task, despite a marked reduction in average neuronal firing rates independent of behavior. These results suggest that the *App* knock-in mutations in A-TG mice induced early AD-like symptoms, including plaque pathology, neuronal hypoactivity, and cognitive impairment before changes in hippocampal neuronal encoding and more severe impairment in behavior.

The mice were trained in two tasks. We used a novel head-fixed air-induced running task in which they learned to run a fixed distance in response to an air stream and to stop running when the air stream terminated. The task was used because it could be presented in a number of configurations. It also allowed for the exquisite control of movement and immobility and response to sensory change events related to hippocampal function (Inayat et al., 2022). The mice were also trained in the Morris place task, a standard assessment for spatial deficits. There were no differences in learning or performance of any of four configurations of the A-TG mice in the motoric aspects of the air-induced running task. The A-TG mice were impaired in learning the place task, but their acquisition of the place task as assessed by a quadrant preference test was not different from C-TG mice. The place learning symptoms observed in the present study appeared milder than those reported in the parent APPki strain at 12 months of age (Saito et al., 2014; Mehla et al., 2019; Jun et al., 2020) and in a variant crossed strain (Takamura et al., 2021). Thus, behaviorally the mice present with symptoms of an early stage of AD in which symptoms of memory impairment occur in the presence of more normal motoric behavior.

The recordings made in the air induced running task showed that neuronal encoding of behavior was similar in the C-TG and A-TG mice. The responsivity of cells to changing features of the task were similar, the proportions of conjunctive and complementary cells was similar, and distance and time encoding was similar. Our finding that time encoding is preserved in A-TG mice is consistent with an earlier report (Takamura et al., 2021). Nevertheless, the seemingly normal distance encoding in A-TG mice is contrasted with disruption of remapping in place cells in APPki mice at 12 months (Jun et al., 2020) and a reduction in the percentage and specificity of place cells in 7-month-old *App^NL-G-F/NL-G-F^*/*Thy1*-G-CaMP7-T2A-DsRed2^+/-^ mice (Takamura et al., 2021). These differences could be due to different stages of disease progression, or to the use of positive reward-based paradigms in previous studies versus the aversive air stimulus paradigm used here. We think that it is unlikely that differences were due to the simplicity of the air-induced running, as we did find neuronal response differences in the four test configurations, as reflected in the comparison of mutual information of Configuration 3 versus Configurations 2 & 1, which suggests that the neurons are sensitive to task parameters. It is possible that a more cognitively demanding task may reveal deficits in neuronal encoding. This is somewhat supported by the observation of mild spatial behavior impairment in the water place task in A-TG mice. It is most likely, however, that the A-TG mice in our experiment were at an early disease stage where the full-blown harmful effects of Aβ pathology at the neuronal level had not yet appeared.

Despite normal neuronal encoding, neurons in A-TG mice were less active than neurons in C-TG mice, both when the mice were running a fixed distance and when they remained still after running. Neuronal hypoactivity may be related to a moderate increase in postsynaptic inhibitory function, which has been described previously in the CA1 region of 6- to 8-month-old APPki mice (Latif-Hernandez et al., 2020). Jun et al., (2020), however, reported no difference in neuronal activity between APPki and *App* wild-type (C57BL/6) mice, whereas (Takamura et al., 2021) found hyperactive neurons independent of behavior in the vicinity of Aβ plaques in 4- to 7-month-old *App^NL-G-F/NL-G-F^*/*Thy1*-G-CaMP7-T2A-DsRed2^+/-^ mice, consistent with observations from calcium imaging of neocortical neurons in earlier transgenic models (Busche et al., 2008). One might argue that the improved calcium sensor used by Takamura et al. (G-CaMP7 vs. GCaMP6s in the present study) may have been more capable of picking up differences in firing rate – although, if this were the case, then the electrophysiological approach used by Jun et al. should have been most sensitive of all, and they observed no difference in average firing rates. The difference between the present study and these previous studies could be related to the proximity of neurons to plaques (which was not assessed in the present study) or, perhaps most likely, to the stage of disease progression. Although both hyper and hypoactivity are reported in transgenic mice with *App* overexpression, behavioral and recording procedures do vary by study (Busche et al., 2012). The present findings suggest that neuronal hypoactivity may be an early manifestation of AD in A-TG mice, nevertheless it may be worthwhile in future studies to make general activity levels of neurons a focus of research with respect to AD symptomology.

The extent of the Aβ pathology seen in A-TG mice at 12 months of age was also less pronounced as compared to similar quantifications done in APPki mice in a previous study in our laboratory (Mehla et al., 2019) and in other studies (Saito et al., 2014; Jun et al., 2020). It is possible that the crossing of strains (APPki with C-TG) may have mitigated the effect of the *App* knock-in mutations, with the C-TG strain perhaps carrying some gene (or genes) that may have triggered mechanisms or pathways protective against the manifestation of Aβ pathology. In humans, genome-wide association studies and meta-analyses have shown that the effect of mutations that increase the risk of developing AD can be mitigated by the presence of protective genes (Dumitrescu et al., 2020; Seto et al., 2021). For instance, the AD risk of the APOE-ε4 gene is reduced in the presence of the Klotho gene. Our study contrasts with a previous study using a similar approach, in which *App^NL-G-F/NL-G-F^*/*Thy1*-G-CaMP7-T2A-DsRed2^+/-^ mice did not show a delay in Aβ pathology and developed neuronal deficits as early as 4 months of age (Takamura et al., 2021). Further genomic and/or proteomic investigations, to identify the genetic mechanisms, with direct behavioral comparisons between strains, might be worthwhile to support.

Overall, this study highlights behavioral and functional relationships of an early stage of AD disease in the A-TG mice. The mice displayed a preserved CA1 network organization measured during a head-fixed task associated with preserved motor function. They also displayed neuronal hypoactivity in the head fixed task and a mild impairment in spatial performance in a free-moving, water-based task. This pattern of results is likely due to the to Aβ deposition throughout the brain including the hippocampal formation. These results are consistent with parallel findings in human AD, in which Aβ plaque load is initially associated with mild memory deficits, but otherwise unimpaired behavior and motor functions. Furthermore, our genetic manipulations for the purposes of calcium imaging may have produced unexpected effects on the AD-like pathophysiology in our mouse model, hinting at unexplored genetic and proteinic mechanisms that may alter the course of AD pathology.

## METHODS

### Animals

All experimental protocols were approved by the University of Lethbridge Animal Welfare Committee and followed the guidelines for the ethical use of animals provided by the Canadian Council on Animal Care. Mice were housed in individually-ventilated Optimice cages (Animal Care Systems) in temperature-controlled rooms (22 °C) maintained on a 12:12 light/dark cycle (lights on during the day). Food and water were provided *ad libitum*. Breeding pairs of *App^NL-G-F/NL-G-F^* knock-in mice [C57BL/6-*App*<tm3(NL-G-F)Tcs>] (Saito et al., 2014) were acquired from RIKEN Center for Brain Science, Japan (RBRC #06344), and a colony of these mice was maintained at the Canadian Centre for Behavioural Neuroscience (CCBN). *Thy1*-GCaMP6s mice [C57BL/6J-Tg(Thy1-GCaMP6s)GP4.3Dkim/J] (Dana et al., 2014) were acquired from the Jackson Laboratory (stock #024275), and a colony of these mice was also maintained at the CCBN. These two strains were crossed to generate a colony of *App^NL-G-F/NL-G-F^*×*Thy1*-GCaMP6s (A-TG) mice. The A-TG mice used for the present experiments were homozygous for the *App^NL-G-F^* gene and hemizygous for the GCaMP6s transgene. Hemizygous *Thy1*-GCaMP6s (C-TG) mice were used as controls. For the *in vivo* calcium imaging experiments, 6 A-TG and 8 C-TG mice were used. For the Morris water task experiment, 12 A-TG and 11 C-TG mice were used. Mice were housed in groups of 2-5 same-sex littermates, unless otherwise noted. Genotyping of all mice was done by polymerase chain reaction, using tissue samples collected by ear notching.

### Morris Water Place Task

The Morris water place task (MWT) was conducted as previously described (Mehla et al., 2019). For 5 days prior to testing, mice were handled briefly once per day. On each test day, mice were acclimatized to the testing room for at least 30 min prior to testing. The test was performed in a circular tank (154 cm in diameter, 50 cm deep), with three distinct visual cues placed outside the tank to facilitate spatial navigation. The tank was filled with water (22 ± 1 °C) to a height of 40 cm. Non-toxic white tempera paint was added to the water to make it opaque. A hidden escape platform (11 cm in diameter) was situated 1 cm below the surface of the water in one of the tank quadrants. The location of the platform remained fixed throughout training. For acquisition training, mice were required to perform 4 trials per day for 8 consecutive days. On a given day, mice were released into the pool from each of the 4 cardinal directions, with the order of the start locations pseudo-randomly determined each day. A trial was terminated when the mouse located the hidden platform, after which the mouse was left on the platform for 10 s before being removed from the tank. If a mouse did not locate the platform within 60 s, a maximum latency score was assigned, and the mouse was guided to the platform. The inter-trial interval was ~20 min. On the 9th day, 24 hours after the completion of acquisition training, a probe test was conducted. This consisted of a single 60 s trial with the platform absent from the tank. Tracking software was used to quantify the following measures for each acquisition training day: average latency to find the platform, average path distance to the platform, and average swim speed. For the probe test, the percentage of time spent in the quadrant where the platform was previously located was measured.

### Two-photon Calcium Imaging in Head-fixed Mice during the Air Induced Running task on a Treadmill

#### Cranial Window Surgery

Between 2.5 and 12.5 months of age, the mice were implanted with chronic cranial windows, using aseptic technique, to allow for calcium imaging of neurons in the CA1 field of the hippocampus. Prior to surgery, the mice were injected with dexamethasone (2 mg/kg, intramuscular), buprenorphine (0.05 mg/kg, subcutaneous), and 0.5 ml (subcutaneous) of 0.9% saline solution containing dextrose (5%) and atropine (3 μg/ml). The mice were anesthetized with isoflurane (4% induction, 1.5% maintenance), and body temperature was maintained at 37 °C using a thermo-regulating heating pad. Lidocaine (7 mg/kg) was injected under the scalp as a local anesthetic, and a section of skin was removed with scissors to expose the skull. A custom-made titanium head-plate was fixed to the skull using cyanoacrylate tissue adhesive and dental cement. Using a drill, a small craniotomy (3 mm diameter) was made over the right hippocampus (center at −2.0 mm AP, 1.8 mm ML, relative to bregma). During drilling, the skull was kept moist and cool by application of brain buffer, and the craniotomy was rinsed and filled with buffer immediately following skull removal. The dura underlying the craniotomy was resected, and a small region of neocortex was removed by aspiration, exposing the hippocampus. Once bleeding had ceased, a cranial window was implanted and attached to the skull using tissue adhesive and dental cement. The window consisted of a metal cylinder (3 mm outer diameter, 1.5 mm long) with a glass coverslip attached to the bottom end using optical adhesive (NOA71, Norland). A rubber ring was attached to the outer edge of the head-plate using dental cement, forming a well to hold the distilled water required for imaging with a water-immersion objective. Post-surgery, the mice received daily injections of meloxicam and enrofloxacin (both 10 mg/kg, subcutaneous) for 2 days. The mice were allowed at least 2 months of recovery from surgery prior to treadmill training and imaging. After surgery, the mice were single housed for the remainder of the experiment.

#### Habituation and training

Habituation to head-fixation was conducted over 3 days. The mice were head-fixed to a stationary post twice per day, with sessions gradually increasing in length from 5 to 30 min. Following habituation, animals were trained to run on a linear treadmill track while head fixed. The treadmill apparatus has previously been described (Mao et al., 2017, 2018). The treadmill was not motorized; all movement of the belt was generated by the mouse. The belt was made from Velcro material (Country Brook). Training was done in the dark, on a blank belt devoid of sensory cues, other than olfactory cues (i.e., urine and feces) deposited by the animal during the training session. A clean belt was used for each mouse, and the belts were washed thoroughly at the end of each training session, to minimize the presence of olfactory cues. The mice were trained for 20-30 min each day to run in response to a continuous, mild air stream applied to the back. To inactivate the air, the animal was required to run a fixed distance on the belt. Following an air-off interval of 15 s, the air would turn on again, initiating the next training trial. The distance the mouse was required to run was increased gradually across training trials, starting each day with 30 cm and increasing to a maximum of 150 cm (i.e., one full lap of the belt). On a given training trial, if the mouse ran the required distance with an average speed greater than 7 cm/s, the required distance on the next trial was incremented by 15 cm. If the mouse failed to achieve the target speed (e.g., due to running too slowly, delayed starting, or repeated starting and stopping), the required distance did not change between trials. The mice were trained until they could rapidly reach the maximum distance of 150 cm and achieve the target average speed of 7 cm/s on >75% of trials across a training session. The mice received a minimum of 3 training days, even if they reached criterion before the third day.

#### Behavioral testing

After the mice reached the training criterion, testing was conducted. In most cases, the mice were tested 1-2 days after the final day of training. In two cases (1 C-TG, 1 A-TG), mice were not tested within this time window. In these cases, the mice were given an additional training session to ensure they still met the training criterion the day before they were subjected to testing. The testing protocol consisted of four different belt configurations, with 10 trials per configuration, and a 1-min rest between configurations. For each trial, mice were required to run one full lap of the belt (150 cm) in response to the air, after which the air was inactivated for a 15 s air-off interval. In Configuration 1, mice ran on a blank belt with no sensory cues. In Configuration 2, four visuo-tactile cues were attached to the belt using Velcro. We describe these cues as visuo-tactile because they had a distinct texture from the belt material, providing a tactile element, and because they were phosphorescent, giving them a visual element. The first cue was situated 28 cm from a fixed reference point on the belt, and the remaining three cues were placed at additional 28 cm intervals. Configuration 3 was the same as Configuration 2, except that a brake was applied during the air-off intervals. For Configuration 4, the cues were removed; thus Configuration 4 was identical to Configuration 1 (blank belt), except the brake was applied during the air-off intervals. The duration of the testing varied depending on the running speed of the mouse, but ranged between 20-30 min.

During testing, activity of hippocampal CA1 neurons was recorded by two-photon calcium imaging. Behavioral data were acquired and synchronized with the imaging data using a data acquisition system (Axon Digidata 1322A). These behavioral data included the distance traveled on the belt (measured by rotation encoders attached to the shafts of the treadmill wheels). A fixed reference point on the belt was also tracked, using reflective tape attached to the underside of the belt, which activated a photosensor once per belt lap. Air onset and offset, as well as the brake, were controlled by a microprocessor board (Arduino Uno).

Mice were excluded from analysis if their average speed across trials for any of the four configurations was less than the training threshold of 7 cm/s. Based on this criterion, 1 C-TG mouse and 1 A-TG mouse were excluded. In addition, a recording from 1 C-TG mouse could not be analyzed due to poor imaging quality resulting from technical difficulties. One C-TG mouse did not complete testing, because it had to be prematurely euthanized due to a broken cranial window. Thus, results from 5 mice per group (A-TG: 3 males, mean age = 417 days; C-TG: 4 males, mean age = 304 days) were compared in the final analysis.

#### Two-photon calcium imaging

A multiphoton microscopy system (Bergamo II, Thorlabs) was used for two-photon calcium imaging. A Ti:Sapphire excitation laser (Coherent) was tuned to 920 nm, with a power of ~100 mW at the sample. Laser scanning was controlled by galvo and resonant scanners. A GaAsP photomultiplier tube was used to detect the green fluorescence signal from GCaMP6s. Samples were imaged through a 16×/0.8 NA water-immersion objective (Nikon). A black fabric cover was wrapped around the objective and head-plate to block contaminating light. Images of hippocampal CA1 pyramidal neurons were captured from a single focal plane ~125 μm beneath the tissue surface, except for one C-TG mouse in which data was collected from two planes with the dorsal plane ~125 μm beneath the tissue surface and the second plane 50 μm deeper. Images were gathered at a frame rate of 29.16 Hz, but the effective frame rate for two-plane imaging was was 9.72 Hz (29.16 ÷ 3, because 1 extra fly-back frame was also captured). A field of view of 418 × 418 μm was captured with a resolution of 512 × 512 pixels.

#### Imaging data processing

Suite2P (Pachitariu et al., 2017) (http://mouseland.github.io/suite2p) was used to register images and to automatically detect regions of interest (ROIs) assumed to represent individual cell bodies. Raw calcium traces were determined for each ROI by averaging the fluorescence signal (F) from all pixels within the ROI and then baseline (Fo) subtracted and normalized to obtain ΔF/F_o_. From the ΔF/Fo calcium traces, deconvolved spike rates were determined using a first order auto-regressive model and constrained nonnegative matrix factorization (Friedrich et al., 2017). Further analysis of the deconvolved traces was conducted using custom scripts written in MATLAB (MathWorks).

To make peri-event raster plots from neuronal activity, a distance bin width of 3 cm and time bin width of 0.3s was used (50 bins for 150 cm belt length and 50 bins for 15 s air-off interval), and the average firing rate was calculated for each time bin. The assessment of whether a cell responded in relation to distance or time was done using a set of independent one-way analysis of variance (ANOVA) tests on the raster plots. In the first ANOVA test, individual bins of a raster were considered individual groups to assess whether mean activity in any bin was significantly different from mean activity in any other bin. In the second ANOVA, adjacent bins (1 and 2, 3 and 4, etc.) were treated as one group, and compared to all other groups. This process was iterated until only two groups remained, representing the first and second halves of all bins in a raster. A cell was deemed responsive if for any one of the 24 independent ANOVA tests *p* < .05. With this agglomerative-hierarchical scheme of analyzing bins of different resolutions with multiple ANOVA tests, a variety of cells with different types of response characteristics were detected. That is, cells with a range of tuning widths were detected, from cells firing in single bins to cells that had distributed firing across bins and distance/time (trials).

Response fidelity (RF) of a neuron for a raster was defined as the percentage of trials in which the firing rate was greater than zero for any bin. The mutual information (MI) between firing rate and distance (or time) was calculated for each raster (PEDR or PETR) using firing rate values from all trials by binning them into four non-overlapping quantiles (Souza et al., 2018) using the following relation:

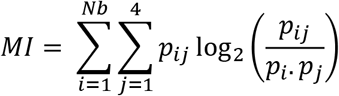

where *p_i_* and *p_j_* are the probabilities of time bin i and firing rate bin *j*, respectively; *p_ij_* is the joint probability between time bin i and firing rate bin *j*; and *N_b_* is the number of distance or time bins (i.e., 50). MI was z-scored (zMI) using a distribution of 500 MI values calculated from randomly shuffled instances of the same raster.

Peri-event distance and time histograms (PEDHs and PETHs, respectively) were obtained by averaging the cellular response in the rasters over trials. Rate vector maps were obtained by plotting the PEDHs and PETHs of all cells after normalizing with the peak response (separately for each cell). The population vector correlation was obtained by finding the Pearson linear correlation coefficient (PLCC) between all pairs of columns of the rate vector map. That is, each pixel shows the PLCC between two columns of a rate vector map (*P_i_* and *P_j_*) given by the following equation:

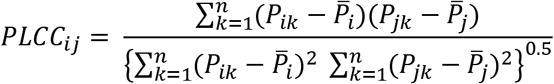

where 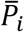 *and* 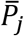 are average values of *P_i_* and *P_j_*, k is the cell number iteration variable, and n is the number of cells. The population vector correlation matrix is hence symmetric across the diagonal. The “Corr” MATLAB function was used to calculate PLCC.

The goodness-of-fit (ordinary R^2^) of a 1D Gaussian fit on each PEDH and PETH was determined after performing least-squares fit with the “fitnlm” MATLAB function using the following equation to estimate a, b, and c:

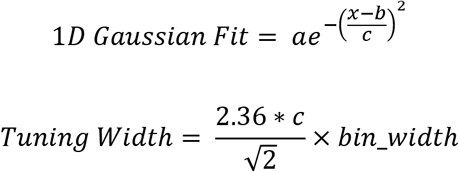

Tuning width was considered as the full width at the half maximum value. For examining the dynamics of activity across configurations, the activity correlation (spatial or temporal) was calculated using the following relation:

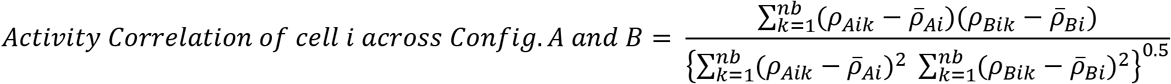

Where *ρ_Ai_* and *ρ_Bi_* are peri-event histograms (average firing rate over trials) for the *i*th cell corresponding to Configurations A and B, respectively, and 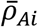 and 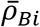 are their average values, k is the bin number iteration variable, and *nb* is the number of bins.

The population vector correlation was calculated using the relation for the PLCC but for the same bins across two Configurations. The rate remapping score was calculated using the following relation:

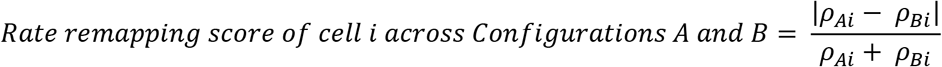

#### Agglomerative hierarchical clustering

As described previously (Inayat et al., 2022), this analysis was performed on the conjunction or complementation matrices (heatmaps) when multiple cell groups were considered. Complementation matrices were split into two heatmaps, symmetric across the diagonal, by using the upper and lower triangular matrices corresponding to Comp1 and Comp2 types of cells before clustering. Based on the conjunction and complementation percentages of each cell group with every other cell group, this analysis found configurations which were similar to each other (i.e., close to each other with respect to their conjunction/complementation relationships with all other cell groups). The analysis was done by first calculating the Euclidian distance for each pair of cell groups, considering two rows of a matrix at a time (using the MATLAB function “pdist”). For example, if a matrix was n × n, there would be 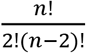 pairs, which can then be arranged into a symmetric n × n matrix. This matrix is usually called a dissimilarity matrix, owing to a smaller Euclidian distance value indicating lesser dissimilarity. The MATLAB function “linkage” was then used to operate on the dissimilarity matrix and define clusters based on the unweighted average distance between clusters. Clusters were then separated (color coded in figures using the MATLAB function “dendrogram”) by using a threshold of 70% of the maximum linkage distance between any two clusters. The quality of clustering was assessed by determining cophenetic correlation (CC) reported with each dendrogram in the figures.

#### Bayesian decoding of distance and time

Decoding of distance and time, with respect to the neuronal population activity, was conducted under a naïve Bayesian paradigm (Zhang et al., 1998). Assuming that the activity of neurons follows an inhomogeneous Poisson process, it can be shown that the maximum *a posteriori* estimate of distance or time is given by:

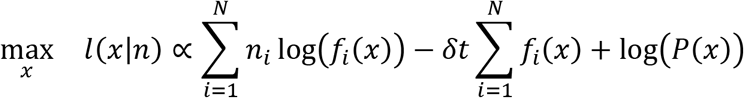

Here, *n_i_* is the deconvolved fluorescence of neuron *i* for a given frame of recording, for a total of *N* simultaneously imaged neurons. *f_i_*(*x*) is the mean activity of neuron *i* at distance or time *x* smoothed using a Gaussian kernel of σ = 10 bins. For both distance and time, this mean activity was discretized over 150 bins. *n_i_* was obtained using a sliding summing window of width *δt* = 2s. For distance decoding, *P(x)* is the probability of occupancy over discrete distance bins *x*. For time decoding, this distribution is flat given that time evolves at a constant rate. Zero firing rates in *f_i_(x)* (which leads to log(0) = –∞) were replaced by a constant penalty term of 2 - 52 (‘eps’ under MATLAB).

Assessment of decoding accuracy was done through leave-one-out cross-validation. Given *k* trials, *k* – 1 trials were used to establish the likelihood and prior probabilities (fi(x) and P(x), respectively). *n_i_* was then extracted from the omitted trial for decoding, and the absolute error between the actual and the decoded distance/time was reported as the decoding error. This process was repeated for all *k* trials.

It is worth noting that the Poisson distribution would not accurately describe the magnitudes of calcium fluorescence (for one, transients in ΔF/F are not discrete events but occur over a continuous positive domain). However, it has been shown previously that such an assumption in practice yields adequately accurate results (Mao et al., 2018; Chang et al., 2020; Esteves et al., 2021). Therefore, as a first-order approximation, the use of this procedure is permissible.

### Histology

To fluorescently label Aβ pathology in the brain, methoxy-X04 (Tocris Bioscience) was administered to the mice by intraperitoneal injection (10 mg/kg; 5 mg/ml in PBS containing 10% DMSO, 45% propylene glycol). Eighteen hours after injection, brains were collected. The mice were deeply anaesthetized with an overdose of sodium pentobarbital, then transcardially perfused with phosphate-buffered saline (PBS) followed by 4% paraformaldehyde (PFA) in PBS. Brains were extracted and post-fixed overnight in 4% PFA in PBS at 4 °C. Brains were then transferred to a sucrose solution (30% sucrose, 0.02% sodium azide in PBS) and stored at 4 °C until sectioning. Brains were cut into 40 μm coronal sections using a freezing, sliding microtome (American Optical, Model #860). Tissue sections were mounted on charged microscope slides (Fisherbrand Superfrost Plus), cover-slipped with Vectashield (Vector Laboratories, H-1000), sealed with nail polish, and stored at 4 °C in the dark until imaging. Images were captured using a slide scanning microscope (NanoZoomer-RS, Hamamatsu).

To quantify Aβ plaque pathology in the hippocampus, images were sampled from every twelfth section of the brain in which the hippocampus was present, for a total of ten images per brain. For each image, both the left and right hippocampi were manually traced. The images were then processed with ilastik (Berg et al., 2019), which was trained to define and count Aβ plaques. Plaque density was calculated for each A-TG brain by dividing the total number of plaques counted by the total area of hippocampus assessed. An experimenter manually verified that no plaques were present in any of the C-TG brains.

### Statistical Analysis

Statistical analysis was conducted using SPSS (version 27) and custom scripts written in MATLAB. For each ANOVA test, the familywise alpha level of each main effect and interaction was controlled at .05. Significant main effects were followed-up using Fisher’s LSD test for comparisons among three conditions, and Bonferroni-corrected pairwise tests for comparisons among four conditions. Significant interactions were followed-up using Bonferroni-corrected simple effects tests (unless otherwise noted), and significant simple effects tests were further followed-up as described for the main effects. For repeated-measures ANOVA tests, the Greenhouse-Geisser correction was applied if the assumption of sphericity was violated (Mauchly’s test: *p* < .05). For these tests, uncorrected degrees of freedom are reported in the text, and the epsilon value is provided. Means presented in the text are reported ± the standard error of the mean (SEM).

## ACKNOWLEDGEMENTS

The authors thank Di Shao for genotyping and animal husbandry, Dr. JianJun Sun for performing surgeries, Hadil Karem and Valérie Lapointe for assistance with histological procedures, and Adam Neumann and Dr. Maurice Needham for technical assistance with two-photon imaging. The authors also thank and acknowledge Drs. Takashi Saito and Takaomi C. Saido (RIKEN Centre for Brain Science, Japan) for providing the *App^NL-G-F/NL-G-F^* knock-in mice used to establish a colony at the CCBN. We also thank Dr. Surjeet Singh for discussions during the preparation of this manuscript.

## AUTHOR CONTRIBUTIONS

All authors participated in the design of this study, with S.I., B.B.M., I.Q.W., and M.H.M as the main contributors. S.I. developed the air induced running task. S.I. and B.B.M. performed the calcium imaging experiments and analyzed data. H.C. performed and S.I. participated in Bayesian decoding analysis. B.B.M performed the Morris water task experiments and analyzed data. S.G.L. performed and B.B.M participated in amyloid beta plaque quantification. S.I., B.B.M, and I.Q.W. wrote the manuscript, which all authors commented on and edited. R.J.S and M.H.M. supervised the study.

